# Petri Net-Based Graphical and Computational Modelling of Biological Systems

**DOI:** 10.1101/047043

**Authors:** Alessandra Livigni, Laura O’Hara, Marta E. Polak, Tim Angus, Lee B. Smith, Tom C. Freeman

## Abstract

In *silico* modelling of biological pathways is a major endeavour of systems biology. Here we present a methodology for construction of pathway models from the literature and other sources using a biologist-friendly graphical modelling system. The pathway notation scheme, called mEPN, is based on the principles of the process diagrams and Petri nets, and facilitates both the graphical representation of complex systems as well as dynamic simulation of their activity. The protocol is divided into four sections: 1) assembly of the pathway in the yEd software package using the mEPN scheme, 2) conversion of the pathway into a computable format, 3) pathway visualisation and *in silico* simulation using the BioLayout *Express*^3D^ software, 4) optimisation of model parameterisation. This method allows reconstruction of any metabolic, signalling and transcriptional pathway as a means of knowledge management, as well as supporting the systems level modelling of their dynamic activity.

## Introduction

The era of molecular biology has resulted in generation of vast amounts of data on biological processes ranging from in-depth studies of one or two molecules and their interactions, to large sets of omics data. These data are currently dissipated in the literature and databases. Those new to a particular biological process or pathway may wish to start researching it through reviewing the primary literature or by reading reviews. However, biological pathways may not have a conventional beginning and end, and the linear prose of a literature review may struggle to present the information in an order that makes it easy to understand as a series of interrelated events. There is also potential for ambiguity as to the exact identity of components due to the frequent use of multiple names for same entity, some or all of which may not be their official names, as dictated by nomenclature committees. Visual representation of biological processes as pathway models can circumvent these issues by presenting the pathway to an end user, allowing them to decide what information is important or indeed missing. Pathway models should logically depict molecular interactions using standardised notation, unambiguously identify the pathway’s components and their transition state, and be easily understood by a biologist. The ultimate goal of a good pathway model is the ability to use it in computational simulations to aid future hypothesis generation. Implementation of a system that fulfils these criteria benefits any scientist working in experimental biology and thus has a large potential user base. To this end, we have combined a proven, user friendly and unambiguous notation system with a Petri net based method of parameterising these models for computational simulation. The use of freely available software with which to construct and run these models makes computational modelling accessible to anyone with a computer and an internet connection.

The modified Edinburgh Pathway Notation (mEPN) scheme was first published in 2008 [1], refined in 2010 [2,3] and is presented here in its current form (Box 1). The scheme consists of entity nodes (which represent biochemical components), process nodes (which represent interactions between the components) and edges, which link entity and process nodes. Pathway maps can be drawn in the free graph editing software yED (yWorks, Tubingen, Germany; www.vworks.com). The bipartite (alternating entity and process node) nature of a pathway model drawn according to the rules of mEPN allows it to be parameterised for computational modelling as a Petri net (a mathematical approach for describing distributed systems, Box 2) by addition of tokens to chosen entry entity nodes in the pathway representing its initial state. When a mEPN model is imported into the open-source software BioLayout *Express*^3D^ a simulation is performed that results in the flow of these tokens through the pathway and generation of quantitative outputs at each entity node that can be visualised as graphs of token flow over time. Altering the simulation parameters can change the flow of tokens, allowing the dynamics of pathways to be modelled under different conditions.

## BOX 1: mEPN Notation 2015

The modified Edinburgh Pathway Notation (mEPN) scheme [3] is a graphical notation system based on the concepts of the process diagram [9], below the glyph library is shown in its current form.

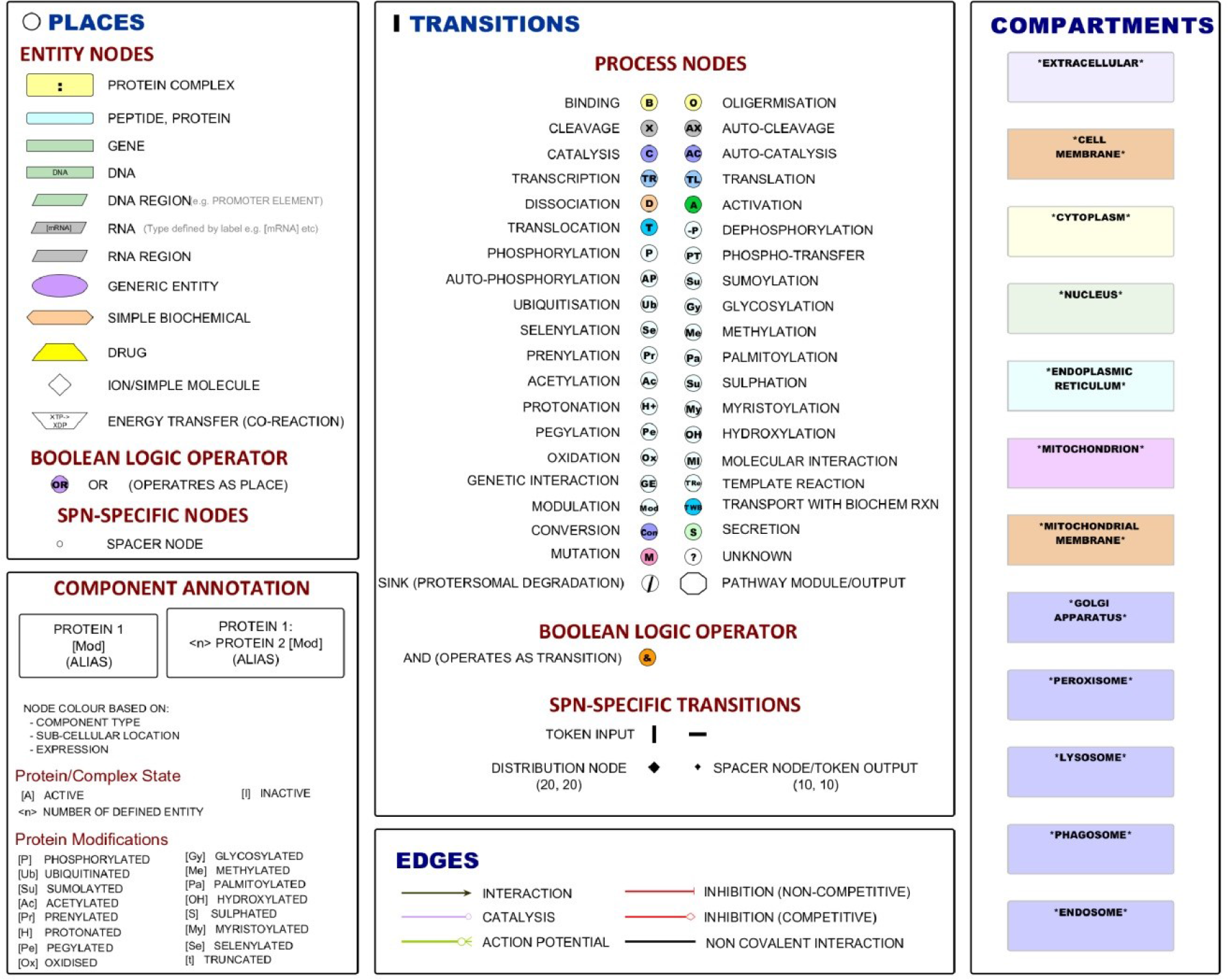

### Pathway components

Nodes are classed as either being ‘entity nodes' (equivalent to Petri net ‘places’) or ‘process nodes' (equivalent to Petri net ‘transitions’) and connected by directional ‘edges’ (equivalent to Petri net ‘arcs’). The scheme consists of twelve different entity nodes (detailed in the top left panel) representing different kinds of molecules present in a biological system. Also included are a Boolean logic ‘OR’ operator node and spacer node (represented as a white circle with a black border) that may be required to maintain bipartite pathway arrangement. All entity nodes are equivalent in Petri net simulations. Included in the scheme are 38 different process nodes (top central panel) representing different types of processes that can occur to or between entities. Additional to these but also acting as transitions are a sink node that is placed at the end of pathway and represents removal of a component from the system, pathway module node that summarises not one process but a pathway of events, Boolean logic ‘AND’ operator node, token input nodes that are placed at the start of a pathway and oriented either horizontally or vertically to fit in with the pathway layout, and a spacer node represented as a black diamond. A larger version of this node can also be used as a distribution node when multiple edges exit from a entity node.

All process nodes are equivalent in Petri net simulations. There are six possible connecting edges (bottom central panel) that represent the nature of the interaction between nodes. The first three (interaction, catalysis and action potential) operate identically within a Petri net and serve to carry tokens between entity and process nodes. The two inhibitor edges act to inhibit the flow of tokens through target process nodes albeit based upon different rules. Finally, the non-covalent interaction edge can be used to depict two separate entities within a complex. This may be a useful thing to do when describing large complexes but these edges do not operate within the context of a Petri net simulation. Pathways can also be compartmentalised into cellular organelles using containers of specific colours (right panel).

## BOX 2: Petri nets to model biological systems

Petri nets are a mathematical approach for describing distributed systems and have been used extensively in the modelling of many different kinds of systems including biological pathways. There are numerous types of Petri net algorithms and software that support modelling using them. However the approach shares a number of common features. They are generally constructed as a directional bipartite networks where nodes are considered to be either ‘places’ or ‘transitions’ connected by ‘arcs’. Places usually represent an entity and by convention are represented as a white circle. Transitions represent interactions between entities or the transition of an entity from one state to another and are usually represented as a black rectangle. Arcs are directional arrows that connect places to transitions and *vice versa*. The availability of a molecule for each process and its abundance can be represented by the initial placement of tokens. The flow of information through the network is represented using tokens that move between places through transitions, following in the direction of the arcs.

### Rules determining flow through transitions

When using a Petri net approach to model a biological system, places represent biological components, such as a cell, protein, complex, mRNA, biochemical etc. Transitions correspond to processes that modify the components in some way, such as phosphorylation, binding etc. The interactions between molecules are depicted by edges, equivalent to arcs in Petri net.

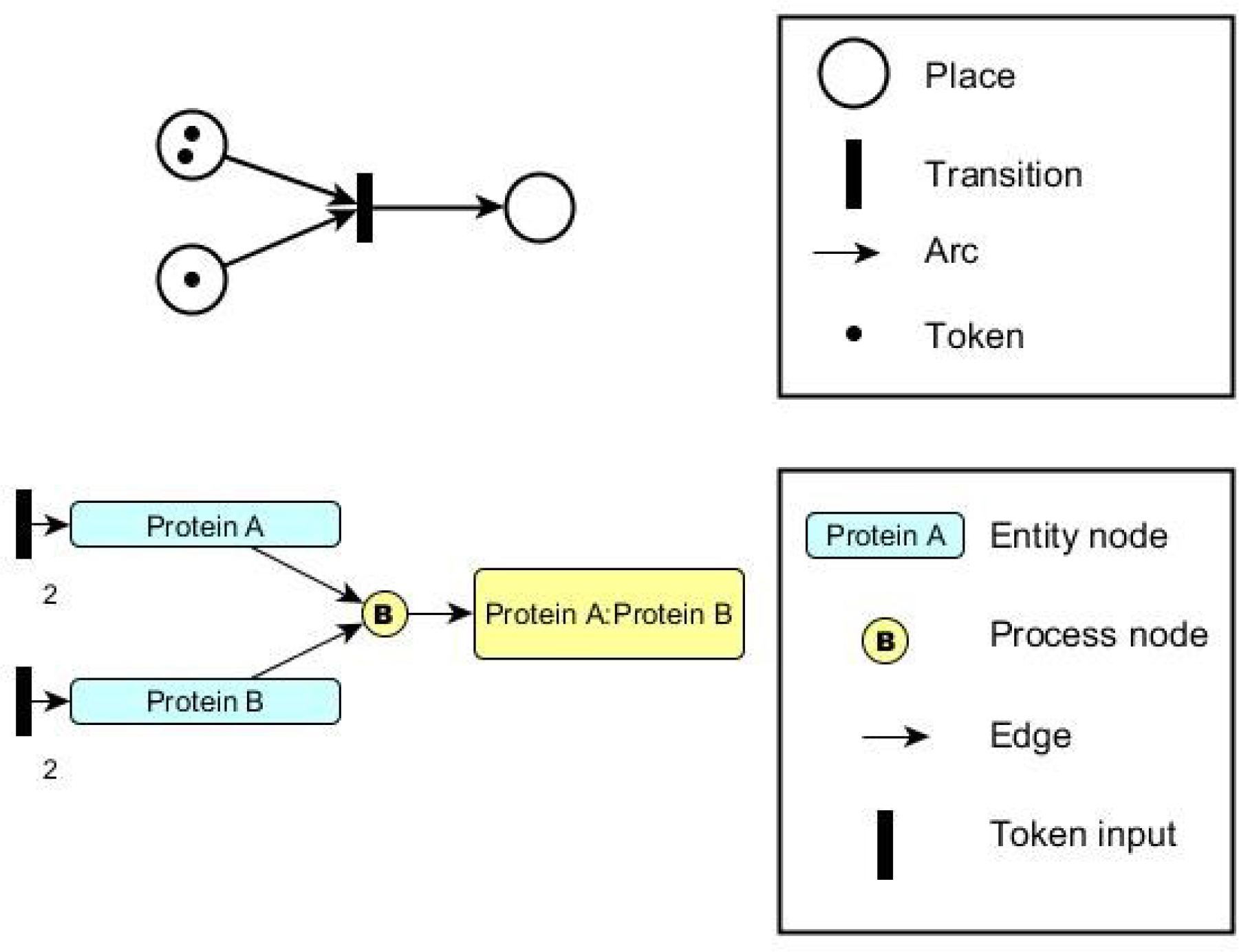

The signal flow is represented by movement of tokens between nodes. The state of each node (network “marking”) is determined by the number of tokens held by it. When a transition is enabled, it may be fired to remove a number of tokens from each input place equal to the weight of the connecting input arc, and create a number of new tokens at each output place equal to the weight of the connecting output arc.

The protocol described here results in dynamic models of biological pathways that will enrich the work of biologists. At their most basic they act as a graphical bibliography, with steps in the pathway hyperlinked to research papers or reports. When fully parameterised and run as dynamic models they can assist with hypothesis generation and identification of gaps in understanding of a particular biological process. *In silico* models can be used in tandem with wet-lab experiments and also have the potential to replace or at least refine them. Construction of dynamic pathway models is a useful tool in a biologist’s skill-set and using mEPN and BioLayout *Express*^3D^ to construct signalling Petri nets means this is now within reach of all biologists, not just those with knowledge of mathematical modelling.

## Experiment Design

First we describe how to construct a graphical model of a biological pathway using the mEPN scheme and then how to convert this graphical model into one that supports computational modelling of the system (steps 1-10). Next, we present how to run simulations and visualize system dynamics (steps 11-17). In the final section (steps 18-23) we describe how to optimise and validate a pathway model. The workflow is shown schematically in Figure 1. To illustrate our approach, we use a relatively simple model of interferon-β signaling, a cytokine pathway which plays an important role innate immunity. To illustrate how to overcome challenges that users may encounter we present a simulation of the interferon-β/SOCS1 feedback response to different doses of interferon-β.

**Figure 1.**
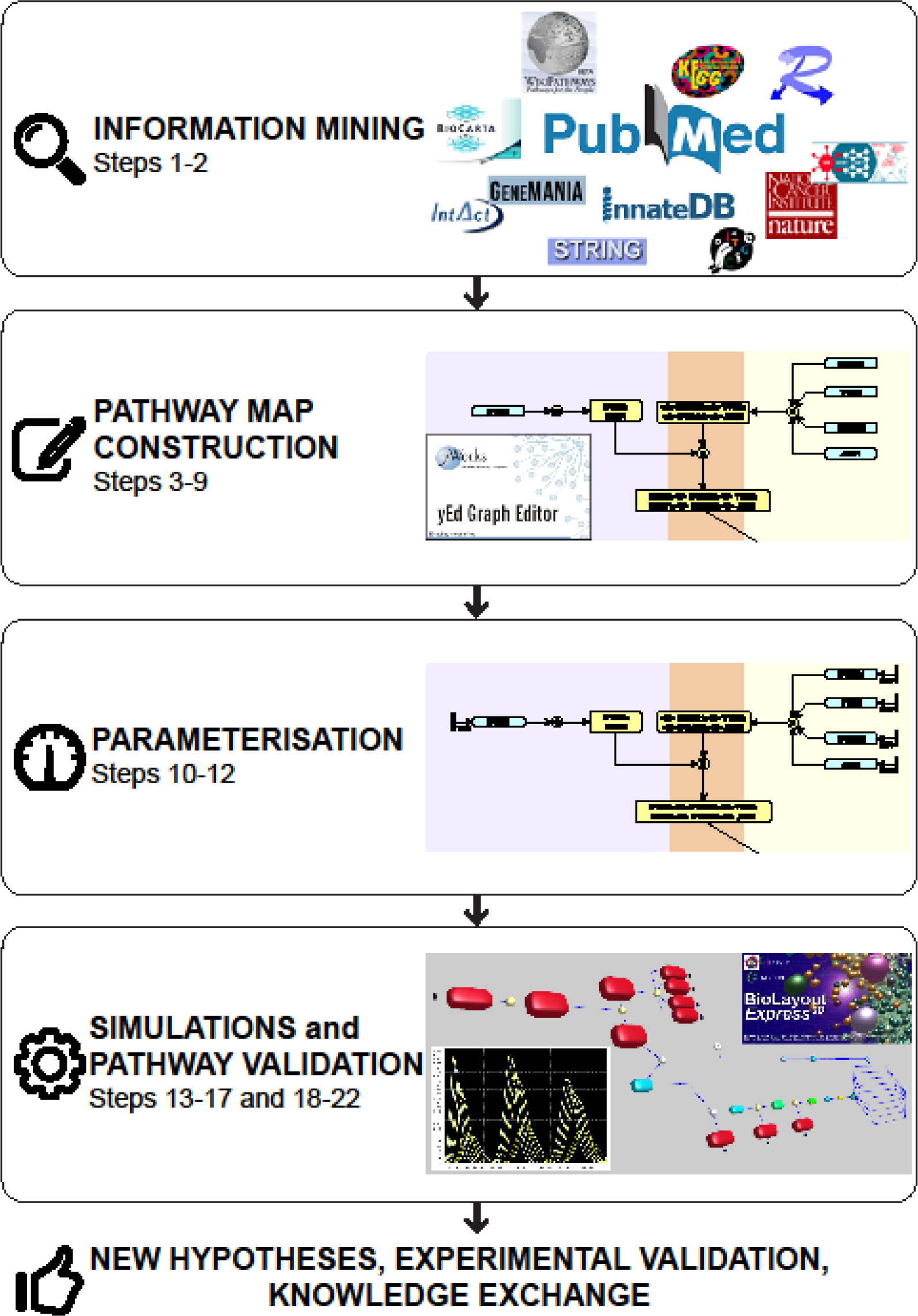

## Materials

### Equipment

A computer with a Windows, Apple Mac or Linux operating system, internet connection and a web browser with JavaScript enabled. The hardware configuration may limit the size of models that can be displayed within yEd, as well as when running pathway simulations within BioLayout *Express*^3D^, where it will influence the speed of simulations and the frame rate for animations of flow. The general rule is that the better hardware, the faster the algorithms will run and smoother the user's interaction with a model. Recommended hardware for optimum performance are as follows:

- main RAM >2Gb
- Dual-core CPU
- NVidia Geforce / Quadro series or ATI equivalent graphics card for advanced visualization with GLSL Shaders
- Monitor capable of displaying at 1,600 × 1,200 resolution Software setup

### Software setup

#### Installation of yEd Graph editor

yEd is a free and intuitive software application that can be used to create high-quality network diagrams, it runs on all major platforms: Windows, Unix/Linux and Mac OS X. Download and install the latest release of the yEd Graph editor from the yWorks (Tubingen, Germany) website www.yworks.com.

#### ♦ Troubleshooting

If you encounter any problems with the installation of yEd contact

#### Installation of BioLayout *Express*^3D^

BioLayout *Express*^3D^ software allows the visualization and analysis of large network graphs in two and three-dimensional space and supports the computational modelling of networks using the signalling Petri net (SPN) algorithm [4]. BioLayout *Express*^3D^ runs on Windows, Mac OS X and Linux platforms. Java SE 6 or 7 is required and can be downloaded from http://www.iava.com/getiava. There are two ways to run BioLayout *Express*^3D^:

- ***Download the BioLayout Express^30^ Installer*** Navigate to **http://www.biolavout.org/download/** and download an installer for Windows (.exe) or Mac OS X (.dmg). For Linux platform use the universal JAR file that may be run without an installer. When BioLayout *Express*^3D^ runs for the first time it creates a preferences file that can be changed and saved at any time from the menu option Tools → Save Preferences. Users can customize many options selecting from the menu bar Tools → General Properties (Shift+P). Further details on the software interface and its customization are available in the BioLayout *Express*^3D^ manual that can be downloaded from http://www.biolavout.org/download/
- ***Launch with Java Web Start*** BioLayout *Express*^3D^ can be run directly from a web browser, on Windows, Mac OS X and Linux platforms, using Java Web Start. The 32-bit version runs from any web browser and will also work on 64-bit systems. The 64-bit build requires 64-bit operating system, web browser and Java.

#### ♦Troubleshooting

If you encounter any problems with the installation of BioLayout *Express*^3D^ contact

## Procedure

### Steps 1 - 7: Pathway model construction

1. **Source information for pathway construction**. A pathway model should aim to provide a comprehensive and reliable view of the current state of knowledge about the system. In order to achieve this one first needs to collect and extract the relevant information about the pathway from the literature, databases and existing diagrams. Possible sources to consult are presented in Table 1. A comprehensive list of databases for data mining can be found on www.pathguide.org. For some pathways, data are available from multiple species and/or cellular systems, therefore users must decide whether to piece together information from heterogeneous models or to restrict their model to a particular system (e.g. human) or developmental stage. To keep track of the data users should create a list spreadsheet that includes: molecules identification (HUGO or Entrez IDs), details about nature of the molecular interactions, the sources (PubMed ID), the quality of evidences and eventual additional information. The use of standard gene/protein names is essential in defining the exact identity of components, especially if models are to be used in the interpretation of omics data where the use of standardised nomenclature systems is standard practice (see BOX 3). Considerable thought should be given to the initial scope of the model, the level of detail to be represented and what the model is to be used for once constructed.

## BOX 3: Component annotation

### Component annotation

For many molecular species multiple names are frequently employed to describe them. This is particularly the case for one of the main components of biological pathways, proteins. Any given protein may be referred to in the literature by a number of different names concurrently. The use of nonstandard nomenclature frequently leads to confusion in written texts and in diagrams the issue is often the same. If the naming of pathway components is not clear then uncertainty arises as to what exactly is being depicted in a diagram and it ends up representing little more than a series of abstract concepts.

Our models have generally been focused on human pathways and we have used standard Human Gene Nomenclature Committee (HGNC) names to label nodes representing genes and proteins. This and related nomenclature systems such as the Mouse Genome Database (MGD) standard, now provide a near complete annotation of all human and mouse genes, and their use in the naming of proteins provides a direct visual link between the identity of the gene and the corresponding protein. Of course not everyone has adopted these naming systems so where other names (alias’) are in common use, these names are often included as part of the node’s label after the official gene symbol in rounded () brackets, but only on its first appearance in the pathway. Use of standard nomenclature also assists in the comparison and overlay of experimental data (which is usually annotated using standard gene nomenclature) onto pathway models. At the present time there are no standard and universally recognized nomenclature systems available for naming certain types of pathway components. For instance protein isoforms tend to be named in an *ad hoc* manner by those who study them and biochemical compounds are known by both their common names or by names that reflect their chemical composition. For instance the IUPAC (www.iupac.org/) provides a standard nomenclature system for organic chemicals, but most names would have little relevance to a biologist. In cases such as these the important thing is to be consistent and where possible to cross reference the components ID to other sources such that the identity of the component depicted, where at all possible, is unambiguous. We have used the excellent ChemSpide resource (www.chemspider.com/) as a reference for the naming biochemical entities although other resources e.g. ChEMBL (www.ebi.ac.uk/chembl/) are potentially equally good. Using the node properties dialogue (F6) nodes may be hyperlinked to external web resources and addition notes to nodes using in the Data tab within yEd.

### Protein state

The particular ‘state’ of an individual protein may determine its functional activity. With mEPN a component’s state is indicated as a text addition to the node label using square [ ] brackets following the component’s name; each modification being placed in separate brackets. The system can be used to describe a wide range of protein modifications from phosphorylation [P], acetylation [Ac], ubiquitination [Ub] etc. and where details of the site of modification are known this may be represented eg. [P@L232] = phosphorylation at leucine 232.

### Protein complexes

Names of the components are given as a concatenation of the proteins belonging to the complex separated by a colon. If the complex is commonly referred to by a generic name this may be shown below the constituent parts in rounded ( ) brackets.

Where a specific protein is present multiple times within a complex, this may be represented by placing the number of times a protein is present within the complex in angle brackets <n>.

A node representing a component may be coloured to impart visually information on component’s type e.g. to differentiate between a protein and a complex.

Similarly other types of pathway components may be represented using a range of shapes and colours-see palette for list of glyphs used to represent different entity types.

A component may only be shown once in any given cellular compartment (in a given state). A component may however alter from one state to another e.g. inactive to active, unbound to bound, in which case both forms are represented as separate entities. To indicate a different state this may be included under the name in square brackets e.g. [A]-active, [P]-phosphorylated.

2. **Download and install the yEd Graph editor**. This is described in Materials/Software Setup. **Download the mEPN palette**. Download the GraphML file containing the latest version of the mEPN glyphs (**Supplementary Data 1**) and load it into the yEd editor by selecting Edit → Manage Palette → Import Selection. This will provide the standard palette of mEPN glyphs that can be selected as required when constructing a pathway model. To display the mEPN symbols palette select from the menu bar Windows → Palette.
3. **Identify the types of pathway information**.Details of a pathway may be divided into the following categories:

- ***Components*** of the pathway: any entity involved in a pathway be it a protein, protein complex, nucleic acids (DNA, RNA), simple biochemical, drug etc.
- ***Interactions*** between the pathway components where one component interacts with or influences the activity of another through its binding, inhibition, catalytic conversion, etc.
- ***Cellular compartments*** are where the components of the pathway reside and interactions take place, such as an organelle (e.g. mitochondrion, nucleus) or a transient compartment (e.g. coated vesicle) etc. As an example, see the list of components present in the interferon-β pathway (Figure 2A).

**Figure 2.**
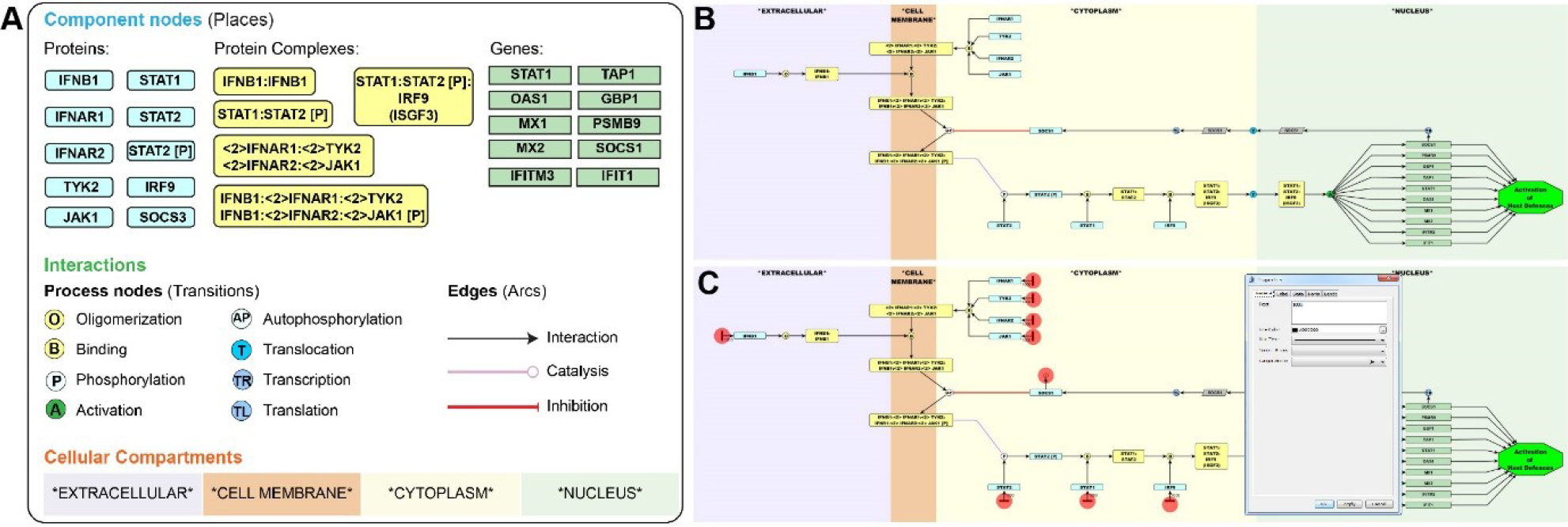
**Construction of a pathway model describing interferon-β signalling**. **(A)** Components of the interferon-β signalling pathway drawn using mEPN notation (see Box 1 for details). The information necessary to construct the interferon-β pathway has been highlighted in the pathway description: components (blue), interactions (green) and cellular compartments (orange). **(B)** interferon-β pathway diagram assembled in yEd software using the components drawn in A. **(C)** Parameterisation of interferon-β pathway by the addition of token inputs and a sink node output on SOCS1 inhibitor node (highlighted by red circles). Also shown is the edge properties dialogue box in yEd where tokens can be added to an input edge.
4. **Commence drawing: add entity nodes**. Molecular components of a pathway are represented using entity nodes which function as the equivalent of a Petri net ‘place’. To add entity nodes to the diagram select and drag the appropriate glyph from the mEPN palette (Box 1). A node’s properties can be edited by selecting it and pressing [F6]. A user can add the component’s name, change the size of the node,record the reference source or insert a brief description about a given component. A hyperlink can also be added to an external site, (for example alink to a PDF of a research paper) which can then be activated by pressing[F8]. A description of the component if available will be shown in a pop-up window when the mouse is placed over the node in yEd. Further details of component annotation are described in Box 1. The sample *Interferon_components.graphml* file can be used in this step (**Supplementary Data 2**).
5. **Draw the interactions between entity nodes**. The nature of an interaction between components may be represented using process nodes and edges.
  - ***Process nodes*** are used to describe the type of interaction that takes place between components and are generally depicted as small round circles with a 1–3 letter code to indicate the type of process e.g. P =phosphorylation, B =binds, X =cleavage etc. A process node may have one or multiple inputs or outputs depending on the nature of the interaction being depicted. As with component (entity) nodes additional information may be added to the node by selecting it and opening the properties dialogue [F6]. Process nodes function as the equivalent of a Petri net ‘transition’. ***Pathway modules*** are a special type of process node. They represent multi-reaction processes or events and are represented using octagons with the name of the process they represent added as a label. They might be used to represent such pathway as signalling cascades, endocytosis, compartment fusion, etc.
  - ***Edges*** are lines that join entity and process nodes. Edges denote the type of interaction (activation, catalysis, inhibition) and their directionality establishes inputs and outputs from entity nodes. Edges function as the equivalent of Petri net ‘arcs’.
  In general and absolutely so when constructing a diagram to be used for computational modelling, nodes comprising a pathway should be arranged as a bipartite graph i.e. entity nodes should be connected exclusively via a process node and vice versa. This structure is the same structure used by Petri nets (places must be connected to transitions) and it is **essential if the model is to be used for simulation experiments**.
6. **Add compartments**. Compartments allow components of the model to be placed in reference to their position within a cell or tissue. They are represented by large nodes generally of a light pastel colour and named according to the compartment they represent, placing the name between asterisks (*compartment name*). The asterisks inform the BioLayout *Express*^3D^ parser to treat these nodes differently: they are displayed as a translucent background to the pathway
and cannot be selected within this tool. If they are labelled as follows *compartment*N* where *N* is a numerical value e.g. 100, when viewed in BioLayout the compartment becomes a 3D container where the N value determines its depth (Z-value). Suggested cellular compartment colours are defined in the mEPN palette. Drag the desired compartment node from the mEPN palette and enlarge it to cover the section of pathway diagram. Move the selected compartment behind the diagram by choosing Edit → Lower selection. The completed pathway should look similar to the interferon-β pathway example in Figure 2B.
7. **Saving and exporting pathway models**. The pathway model should now represent a diagrammatic version of known events and should be saved in the GraphML file format choosing File → Save. A model can also be exported as an image in PNG or JPG format, as PDF or as HTML by selecting File → Export.

### Steps 8 - 10: Conversion of a graphical model into a computation model

8. **Check token flow through the model**. Care must be taken to ensure that the model conforms to the bipartite structure required by the SPN algorithm (as described in Box 2). When necessary, spacer nodes (Box 1) representing either a place or a transition can be added in order to preserve the bipartite structure of the model. Whilst the various entity and process nodes defined by the mEPN scheme may appear different, they are all considered to be either places or transitions, respectively, in terms of a Petri net. All entity nodes can carry tokens and all process nodes function identically as transitions within a Petri net simulation i.e. the rules governing token flow through them are the same regardless of the type of entity or process they represent.
9. **Set the initial parameters**. In order to convert a graphical representation of a pathway diagram into a computational model, the initial state of the system must be defined through a process of parameterisation. In signalling Petri nets, the primary way of achieving this is by the placement of tokens. Tokens represent the amount or activity of a pathway component (e.g. the level of gene/protein expression or accumulation of a product/substrate) and are generally added to entity nodes at the beginning of network i.e. nodes that have no ‘parents’ (upstream connections). The number of tokens assigned to a node can be derived from an experimental value (e.g. gene or protein expression level) or have an estimated value. For an example of where tokens should be placed in the interferon-β pathway see Figure 2C, marked by red circles. In order to define the initial state of a component, an input node (depicted as a black rectangle which functions as a transition node), is placed upstream of the component to be parameterised and connected to it with an edge. The number of tokens to be added to the node is defined by selecting the edge between the input node and component. The edge properties dialogue [F6] is opened and the desired number of tokens is typed into the edge name (text) box (Figure 2C). Token values can range from 0 to millions but in essence represent the relative amount or activity of a given component under initial conditions. Ideally the initial parameters should be set with reference to some experimental data providing information on the relative initial concentrations of the pathway components where known. However in the initial stages of pathway parameterisation it is often sufficient to place an arbitrary number of tokens on component just to check the connectivity between inputs and outputs is not compromised in any way. The dynamic behaviour of the pathway is then simulated by the movement of the tokens between entity nodes (token flow) by the algorithm executed within BioLayout *Express*^3D^.
10. Preparation for simulation experiment. Once a model has been parameterised the ‘SPN compatible’ version of the pathway can be saved as in the GraphML file format choosing File → Save. GraphML files can be loaded directly into BioLayout *Express*^3D^. A parser within the tool translates the mEPN nodes into their 3D equivalent shapes such that they can now be visualised within the tool’s 3D environment. It is also designed to differentiate which nodes in the diagrams act as Petri net ‘places’ and which are ‘transitions’, and the parameterisation markings that define token input and flow. The tool is also able to perform stochastic flow simulations using a modified version of SPN algorithm.

**Steps 11-17: Visualization of pathway and running simulations using BioLayout *Express*^3D^**

11. **Install BioLayout Express^3D^ and open pathway models**. Install BioLayout *Express*^3D^ as described in Materials/Software Setup and open the program. When a pathway model is created in yEd and saved as a GraphML file is loaded into BioLayout *Express*^3D^ a dialogue will appear that asks “This looks like a Signalling Petri (SPN) pathway. Would you like to run a SPN simulation now?” (Figure 3A). When the answer ‘Yes’ is clicked the SPN simulation dialogue appears (Figure 3B). The dialogue can also be selected from main menu under the Simulation menu or by pressing the “RUN SPN” button on the sidebar.

The sample *Interferon_SPN_ready_model. graphml* file can be used in steps 12-17 (**Supplementary Data 3**).

**Figure 3.**
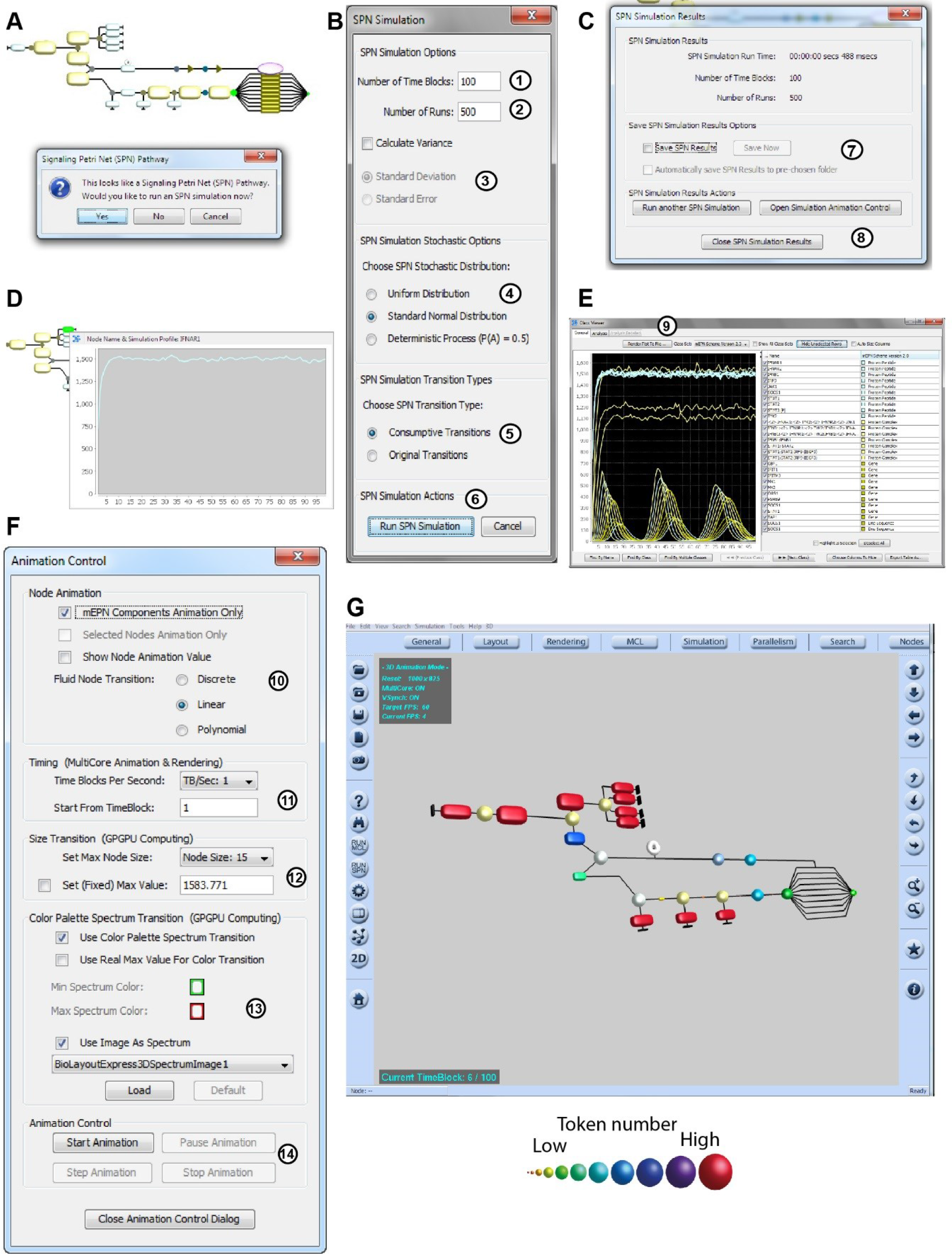
**Visualisation of token flow**. **(A)** When a model is loaded into BioLayout *Express*^3D^ it is displayed in a 3D environment using for the most part 3D equivalents to the 2D node glyphs as rendered in yEd. The software automatically recognizes a diagram as having been parameterised for computational modelling (based on the presence of process nodes as defined in the notation system) and prompts users to run a SPN simulation. **(B)** In the SPN simulation dialogue users can set constrains on how to run the SPN simulation algorithm. (1) Define the number of time blocks in a simulation; (2) Define the number of runs in a simulation; (3) calculate variance in the stochasticity of flow between runs; (4) define the nature of the stochastic flow of tokens; (5) define the rules governing token flow at transitions; (6) the user may then run the SPN simulation. **(C)** SPN simulation results dialogue box summarises the SPN simulation (7) SPN results may be saved before the user chooses to (8) run the simulation again, close the dialogue box or proceed to animate the simulation. **(D)** After a simulation has been run token accumulation at specific nodes can be visualised by placing the cursor over the node. **(E)** The flow of tokens across one or a number of selected nodes can be plotted using the Class Viewer. Graphical plots of token flow over the time course of the experiment for selected nodes. The name and class of selected nodes is also displayed, and below a range of options are available for node selection and data export. (9) The token plot can be saved as a.png or.jpg image file. **(F)** Animation control dialogue. (10) Options for the type of nodes to be animated and interpolation of flow between time bocks; (11) speed of animation (time blocks per second); (12) node size at maximum flow; (13) colour palette selection for highlight change in token number; (14) animation control: start, pause, step, stop. **(G)** BioLayout *Express*^3D^ can produce animations of token flow through the model across time blocks. Node size will increase/decrease depending on the number of tokens passing through them and the colour of the node will also change according to a predefined spectrum of colours.

12. **Run a pathway simulation**.When running a pathway simulation a number of user-defined options are available.

- **SPN Simulation Options**

- *Number of Time Blocks*: a time block is a when all transitions are fired exactly once in random order. A series of time blocks is referred to as a run (Figure 3B1).
- *Number of Runs*: a simulation maybe repeated multiple times (runs) and the outcomes from individual runs are averaged to calculate the mean number of tokens present on a given node across the time blocks (Figure 3B2). Check the ‘Calculate Variance’ box if required and pick either standard deviation or standard error (Figure 3B3).

- ***SPN Simulation Stochastic Options***: It is possible to change the nature of stochasticity in the distribution of token flow using the following options (Figure 3B4).
- *Uniform Distribution*: each time a transition is fired, an entirely random number of tokens (between zero and the maximum number of tokens on the place) are moved from an input place to the output place (assuming there no other inputs on the transition which may influence flow). This mode is as originally described by Ruths *et al*; [4] in their description of the SPN algorithm.
- *Standard Normal*: each time a transition is fired, the number of tokens moved between input and end places will be randomly chosen from a standard normal distribution around 50% of the number of tokens on the input place.
- *Deterministic*: moves exactly half of the tokens from input place to the output place each time a transition is fired.
- ***SPN Simulation Transition Type***: Flow between places is via transition nodes which all operate using the same set of rules governing token flow. We have introduced two options as to how they operate with respect to token accumulation (Figure 3B5).
- Consumptive Transitions: tokens are consumed from place nodes irrespective of whether the transition is ‘open’ or not i.e. if there are two inputs into a transition and one has tokens and the other does not, tokens will still be lost from the input place with tokens as if flow were unrestricted.
- Original Transitions: tokens accumulate on input nodes where flow from them is blocked i.e. if there are two inputs into a transition and one has tokens and the other does not, tokens will not be lost from the input with tokens. This mode was as originally described by Ruths *et al*; [4].

After selecting the options for running a flow simulation, press the ‘Run the Simulation’ button to initiate the computation of the SPN algorithm (Figure 3B6). The time it takes to run a computation depends on the number of time blocks/runs, the size of the pathway model and hardware on which the simulation is run. However, for most small to medium size pathways and hardware configurations, the time taken is usually a few seconds or less.

13. **Save the results**. Once the SPN simulation algorithm has finished, a Simulation Results dialogue appears (Figure 3C). At this point the results, token level per node per time block can be saved as a.txt or.spn file by ticking the “Save SPN Results” box or pressing ALT+S (Figure 3C7). Saved simulation results files can be loaded pressing ALT+L back in the main window. Results files can also be viewed as a spreadsheet e.g. in Excel. An mEPN model can also be exported as a Systems Biology Graphical Notation (SBGN) [5] diagram via File -> Export -> SBGN file. This can be opened in any SBGN compliant software e.g. VANTED with SBGN-ED add-on [6].
14. **Visualize token flow as node output graphs**. To visualize the simulation results for a selected entity/place node, close the SPN simulation results dialogue box (Figure 3C8), position the cursor over the node of interest and a pop-up window will appear showing the token flow associated with that node (Figure 3D). In order to view and compare token flow in multiple nodes, select the nodes of interest by pressing Shift+left mouse button and dragging the select window over the nodes of interest or by pressing the ALT+left mouse button to select multiple nodes. The corresponding flow graphs can be viewed using the Class Viewer by pressing CTRL+C or the button with Figure 3E10).
15. the icon on left menu bar (Figure 3E). A range of options are available within BioLayout to adjust graph appearance. Pressing the ‘Render Plot to File button’ on the top of the Class viewer window the graph can be saved as *.jpg* or *.png* image file (Figure 3E9). To visualise the SPN output as an animation, open the Simulation Animation Control (ALT+A) window when the simulation has finished (Figure 3C8).
16. **Control visualization of token flow** in the Simulation Animation Control window (Figure 3F).

- **Node Animation** Provides options to choose which nodes are animated and the type of animated transition that takes place between the node value associated with one time block and the next (F
- **Timing** (Figure 3E11):
- Time blocks per Second: controls how many time block values are displayed per second of animation and therefore the speed of the animation
- Start from time block: controls the time block from which the animation begins.
- **Size Transition (Figure 3E12)**:
- **Set max node size**: option controls the maximum size of nodes during the visualization of token flow.
- Set (fixed) node value: the number of tokens required at which point the maximum node size and colour is represented. The default value for the ‘Max value’ is determined by the maximum number of tokens that accumulate at any given node during a simulation. It is often the case that some nodes accumulate tokens much in excess others e.g. when output is blocked. This can result in the majority of nodes seemingly to change little in size or colour during a simulation. By manually changing this value and checking the associated box, a user-defined value will be set and maintained for subsequent runs.
- **Colour Palette Spectrum Transition (Figure 3E13)**: This dialogue provides options to control the changes in colour as node token values change. A range of colour spectra are provided or can be user specified.

17. **Visualize token flow as an animation**. When the animation parameters have been set, select ‘Start Animation’ (Figure 3F14) to watch tokens flow through your model (Figure 3G and Supplementary Data 4).

### Steps 18 - 24: Model optimisation and parameterisation

18. **Pathway Layout**. A pathway model should ideally be a compact and easy to follow. Organizing the pathway model into separate 'modules’ based on connectivity facilitates reading and allows for network expansion as new data become available. Further information on layout optimisation can be found in Box 4. In practice model optimisation is normally an iterative process often requiring a degree of trial and error in both the layout of the diagram and in exploring its dynamic properties. It takes time and considerable effort to layout pathway diagrams, but the effort is worth it.
19. **Error checking**. Errors in a diagram’s structure, predominately failure to adhere to the strict requirement for the maintenance of a bipartite graph or improper logic, can lead to bottlenecks in token flow. When the bipartite graph structure is not maintained e.g. an entity node is directly linked to another, the SPN algorithm cannot operate and tokens will accumulate on the node upstream of the issue and zero tokens will be passed on to subsequent nodes. To avoid bottlenecks users should check the reactions preceding any node whose token output is zero (Figure 4A) and correct mistakes as necessary.

## BOX 4: Layout Optimisation

There is a part of pathway modelling that could be considered art, or at least creative cartography. When starting a diagram the number of components is small and visual comprehension of the system of nodes and edges relatively easy. However, this situation soon changes as a diagram grows and one of the greatest challenges is to render the inherently complex connections between components of a network model in a human readable form. The primary tool for this job is the careful placement of nodes and edges in the network layout. There are large number layout algorithms available for network visualisation but unfortunately none come close to the results achievable by as skilled human curator. Of course certain rules can be applied to this process:

- Models should be constructed where possible along a horizontal or vertical axis, arranged top down or left to right in the direction of information flow.
- Nodes should be evenly spaced and aligned along the chosen axis of layout.
- Crossing over of edges should be kept to a minimum and changes in edges direction should be avoided when possible. When multiple edges run parallel to each other it is important to keep the lines straight to maintain an easy-to-follow diagram.
- Space can be organised effectively by structuring sub-pathways into modules.
- Modules of the pathway should be arranged so that the connected glyphs are in close proximity, to minimise tangling of connective edges.
- Hierarchical relationships between components should be shown in the layout of interactions. In order to do this an orientation of pathway flow is chosen (e.g. left to right or top to bottom) and should be maintained throughout the diagram where possible.

However, each diagram is essentially unique and each comes with its own challenges. There is no one solution that fits all models and as diagrams so must adapt the layout to take in new components and concepts.

**Figure 4.**
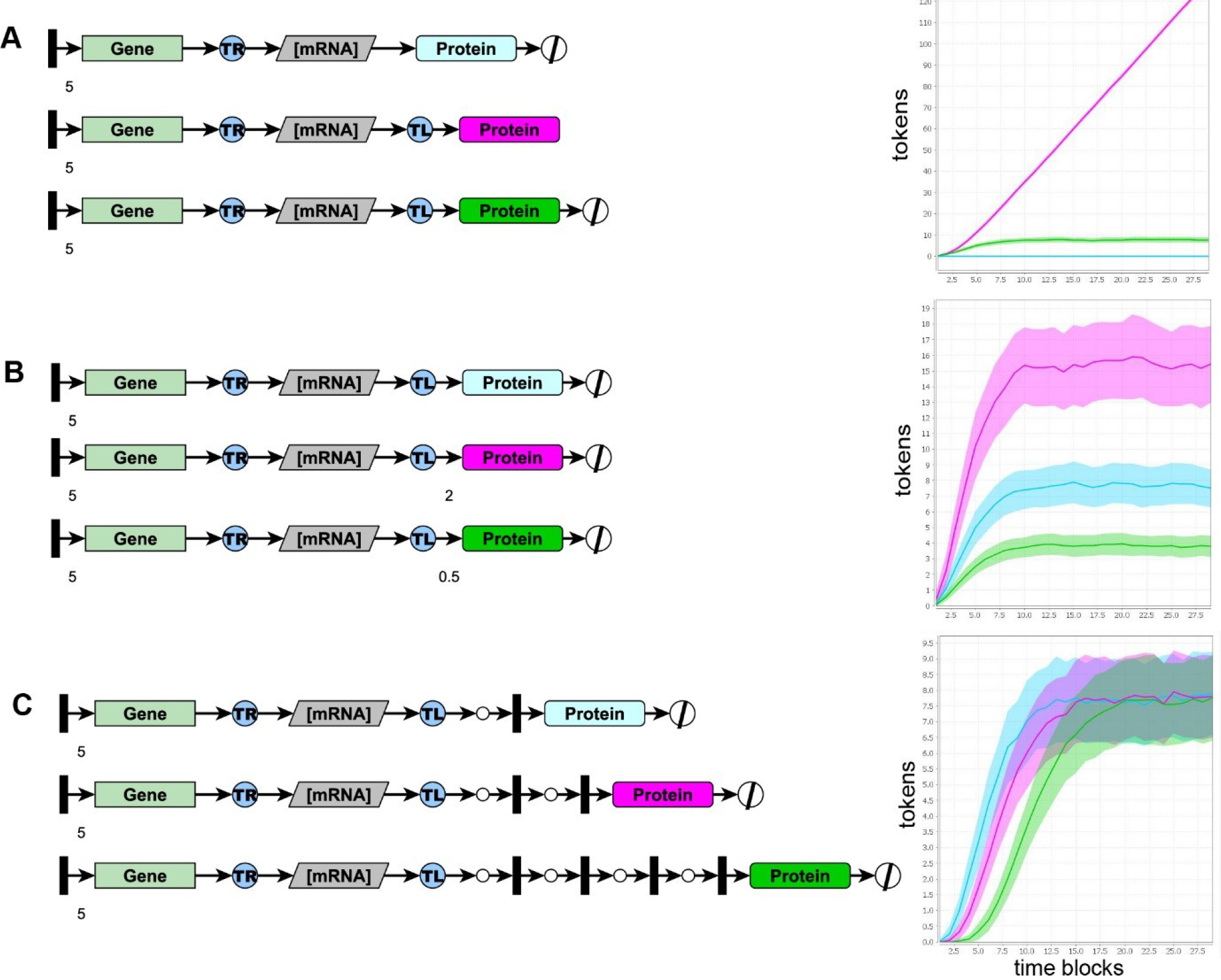
Influencing token flow along a linear pathway. (A) Visualization of blocked reactions. Two entity nodes (mRNA and capped mRNA) have been connected without using a process node. This leads to a bottleneck with exponential accumulation of tokens on the mRNA (yellow) and zero output tokens on the protein (blue). (B) Modifying edges’ weight. The addition of a value to the output edge increases the token output simulating a boost in translation. (C) Modelling time. The graphs show the simulation run using linear networks containing delays of different length. Input tokens are fired simultaneously in each of the networks but the output tokens accumulate at different time blocks. The delay is proportional to the number of transition steps introduced in the network. Simulations in A, B and C: 500 runs, 20 time blocks, stochasticity setting Normal Standard distribution.

20. **End of the line**. At the end of any pathway is potentially a next, but as yet undefined, event or outcome. Without an output a component (place) at the end of a line of flow will simply accumulate tokens as there is nowhere for them to flow on to. For instance, a pathway may end at a gene which is shown to be up-regulated upon activation of the upstream signalling components but it will never turn off even if its activation stops. One approach to stop tokens accumulating at such sites is to place a ‘pathway node’ (a process node defined as a transition), after the gene indicating what the transcribed protein may go on to do. Tokens will then be lost from the system and not accumulate on the entity node representing the gene. Some branches may end in a ‘sink’ process node signifying that a component is removed from the system. Sinks are often used to indicate the removal of a protein by proteosomal degradation.
21. **Amplifying or reducing token flow at specific sites**. Users can simulate the amplification or reduction of signal at specific sites in the network. This is achieved by adding a numeric value to a transition-to-place edge. Select the edge, press [F6] and write a number in the Text field. During the simulation tokens will be transferred according to an edge’s weight. When a transition fires, the number of tokens produced in the output places will correspond to the number of input tokens multiplied by the weight of the output edge e.g. an edge weight of 2 will result in a doubling in the number of input tokens, where as an edge weight of 0.5 will halve the number of tokens going forward. For example, one can amplify the effective rate of translation of specific transcripts by modifying the weight of the output edge from a translation node (Figure 4B).
22. **Variable token input**. Users can also simulate variation in the level of an input signal at different time blocks of a run by changing the token number assigned to an input edge (as described in Step 9) to the following notation: a-b,c;d-e,f where ‘a-b’ are the first and last time blocks that you would like the number of tokens ‘c’ to be added to the model and ‘d-e’ are the first and last time blocks that you would like the number of tokens ‘f to be added to the model. This allows modelling of a system before and after a stimulus, or when a stimulus is transient or delayed.
23. **Modelling time**. Time in Petri nets is measured in abstract units called time blocks. When constructing a model it is useful to consider how many experimental seconds, minutes, or hours correspond to a time block. Timing depends on the network topology and given that flow occurs over consecutive time blocks the further away a node is from the start of flow the longer it will take to accumulate tokens. A transition will pass on upstream tokens only if all the input places contain tokens. So the firing time of some reactions can be controlled by the availability of specific components. As an example, interferon-p responsive genes can only be activated after the cell is subject to interferon stimulation. Some processes have multiple intermediate steps which delay the accumulation of the final components (output nodes). For instance it is possible to represent the translation of mRNA to protein as a single transition ignoring the fact that it represents a complex process comprised of a significant number of individual steps that in reality take time to complete. To simulate such time delays users can create a linear network that alternates transitions with spacer nodes multiple times. When tokens are passed through such a linear network the number of output tokens corresponds to the input but the time taken for tokens to reach the end is proportional to the number of spacer nodes (Figure 4C).
24. **Modelling negative feedback loops**. Such regulatory systems are common and essential in biological systems limiting the degree to which a given process is activated. The network motif is common to many systems and involves the activation of a pathway component that goes on to inhibit an earlier step in the process. The inhibition of the interferon-p receptor by SOCS1 whose genes is activated by the interferon signalling pathway represents such a negative feedback loop (Figure 2). In order to represent an inhibitory activity such as this using the mEPN scheme, an inhibitor edge is placed from the inhibitor molecule to process node representing the step that is inhibited. During a simulation tokens will not be lost through an inhibitor edge. With no output an inhibitory node will accumulate tokens and will irrevocably block the transition to which it is connected. If a sink node is placed on an inhibitor molecule it serves to give the inhibitory molecule a ‘half-life’ (in practice any process node will serve the same purpose, but use of the sink node helps visually define the process involved). In the absence of further input into the inhibitor node, such as during the ‘off phase’ of negative feedback system, tokens will now be lost from the inhibitor. The result is that its inhibitory effect will lessen and the blocked transition will eventually open and tokens may flow again through it. In presence of a constant input this can cause token flow in negative feedback systems to oscillate. There are two types of inhibitor edge included in the notation scheme that perform differently in the modelling environment; the non-competitive inhibitor edge (red with perpendicular bar at end) and a competitive inhibition edge (red with open diamond end). The non-competitive inhibitor edge completely blocks token flow through the target transmission if any tokens are present on the inhibitor node. In contrast the competitive inhibitor edge works by deducting the number of tokens residing on the inhibitor away from the number of tokens flowing through the target transition. As can be seen in Figures 5A and B the effect of these two edges are markedly different. The behaviour of negative feedback systems is not only dependent on the type of inhibitor edge used but also the distance between the input of tokens and the inhibitory step, the great the distance the more time tokens are able to accumulate in the system and time taken between the opening and closing of the inhibited transition i.e. the longer the wavelength and an increase in wave height of the oscillating signal (Figure 5C and D). Other factors that can affect the oscillatory behaviour of feedback loop are the number of inhibitors acting on the pathway, their position and assumptions about the stochasticity of token flow through the system.

**Figure 5.**
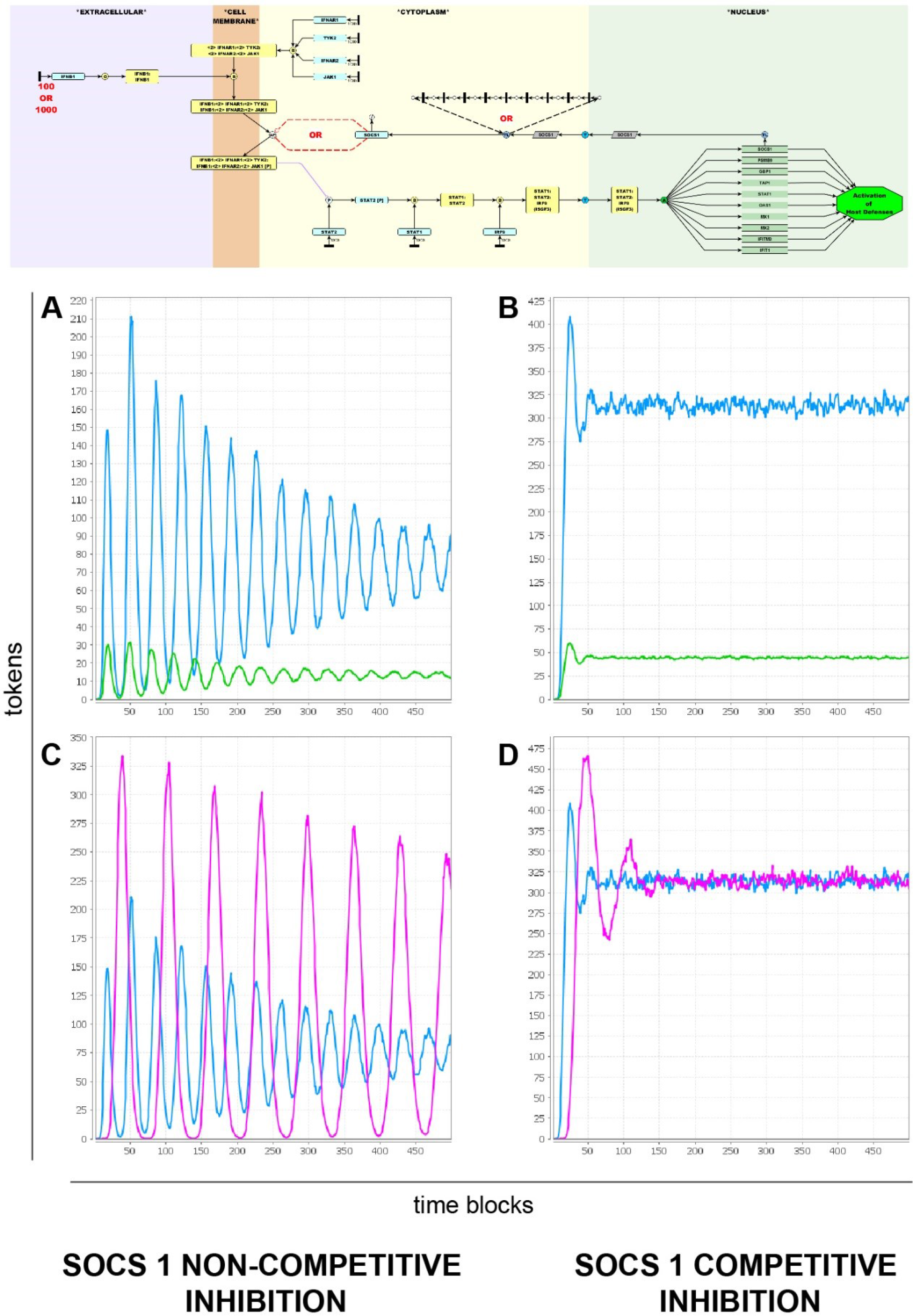
**Parameterization of the interferon pathway**. Graphs show token accumulation on the activated the transcription complex ISGF3 (marked by red X) following simulation of **(A)** a high-(1000 input tokens, blue line) or low-dose (100 tokens, green line) of interferon-β with SOCS1 acting as a non-competitive or **(B)** competitive inhibitor of the activated receptor (dashed red edges). **(C)** Oscillatory activity of the SOCS1 feedback loop with (magenta line) or without (blue line) a delay introduced between the transcription and translation of SOCS1 (dashed black edges), again with SOCS1 acting as a non-competitive or **(D)** competitive inhibitor of the activated receptor. In all cases an input of 1000 tokens was assumed for interferon-β. Simulation: 500 runs, 300 time blocks, stochasticity setting Normal Standard distribution.

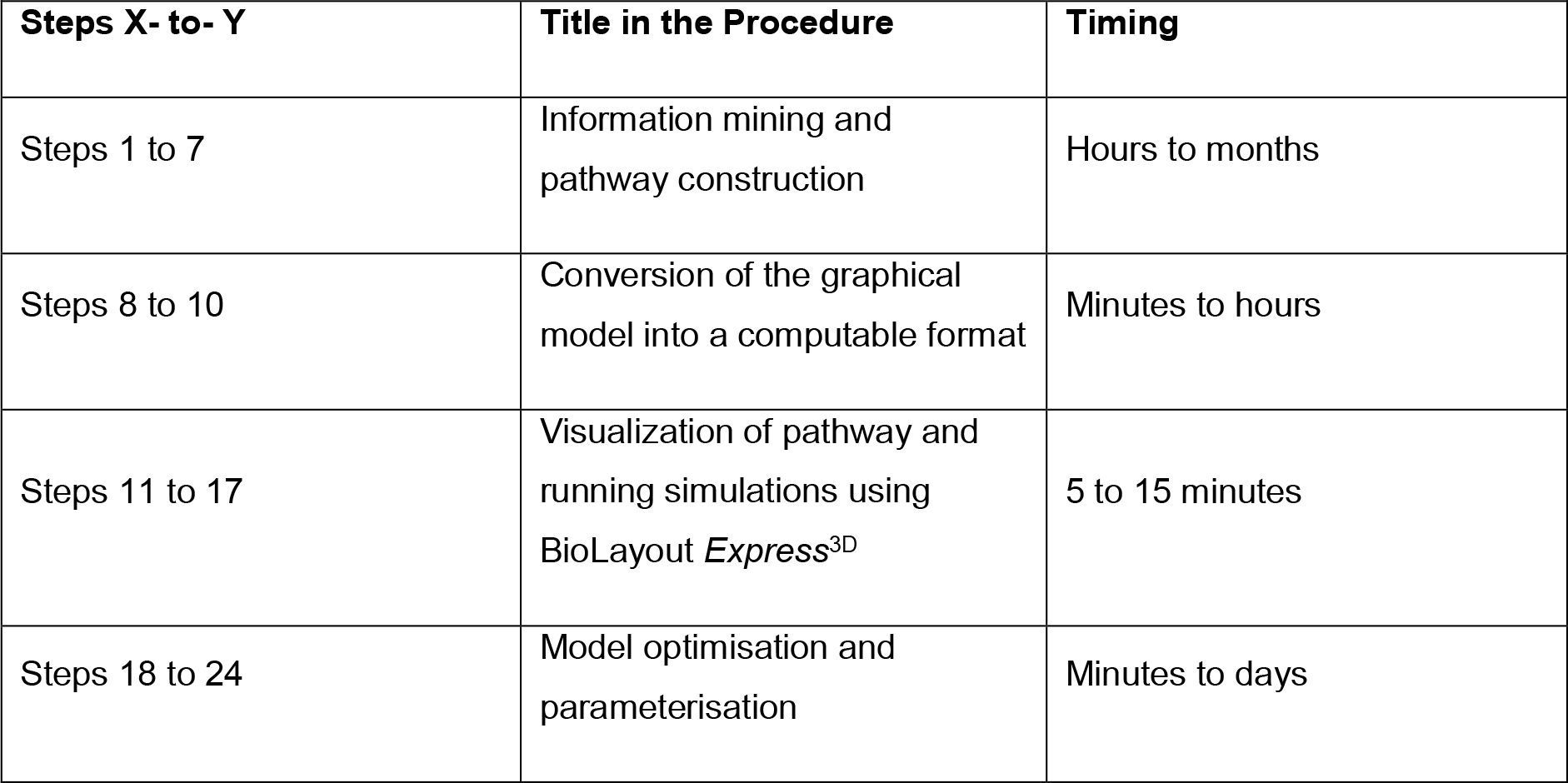
Timing

Timing
| Steps X- to- Y | Title in the Procedure | Timing |
| --- | --- | --- |
| Steps 1 to 7 | Information mining and pathway construction | Hours to months |
| Steps 8 to 10 | Conversion of the graphical model into a computable format | Minutes to hours |
| Steps 11 to 17 | Visualization of pathway and running simulations using BioLayout *Express*<sup>3D</sup> | 5 to 15 minutes |
| Steps 18 to 24 | Model optimisation and parameterisation | Minutes to days |

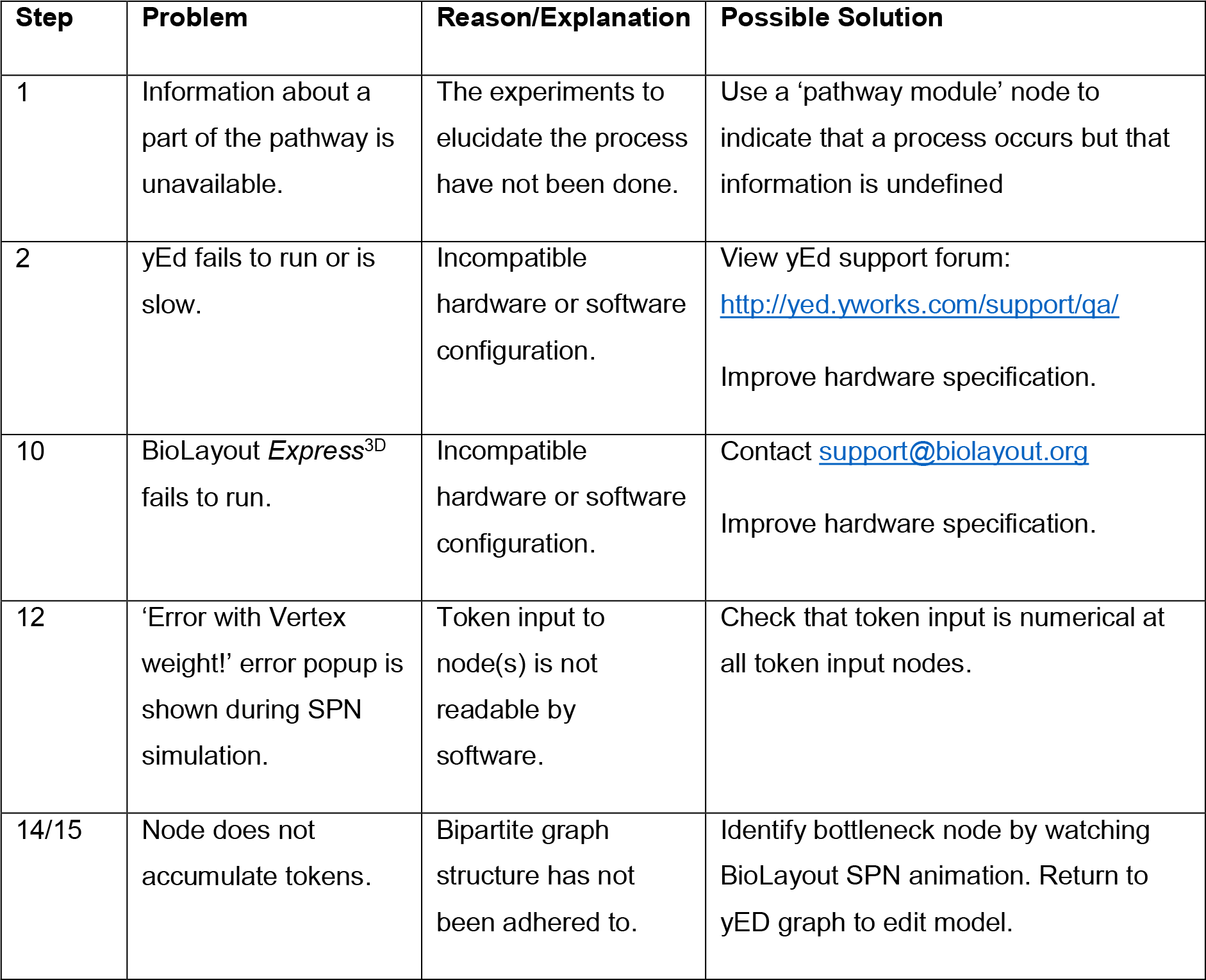
TROUBLESHOOTING

## Anticipated Results

This protocol describes the generation of pathway models using the mEPN language for graphically representing biological systems. mEPN models can be used both as a resource to store and display what is known about a pathway and in turn converted to a computational model by setting parameters to define the initial state of the pathway. Using the sample interferon-β components GraphML file (**Supplementary Data 2**) a pathway model can be produced that represents events from the binding of interferon-p to its receptor and the signalling pathway leading the activation of target genes. The resulting model can be parameterised adding tokens to obtain a SPN-ready model, as provided in the sample interferon-β pathway GraphML file (**Supplementary Data 3**). This is a relatively small diagram, to view and run larger pathway models go to:www.virtuallyimmune.com.

The BioLayout *Express*^30^ software is fully compatible with the mEPN notation and can be used to for pathway visualization and to perform stochastic flow simulations using a modified version of SPN algorithm [4]. Running simulations provides insights into the dynamic behaviour of the system by enabling users to visualize the signal flow within the network. The signal flow is simulated by the accumulation of tokens at entity nodes (places) and can be visualized in 2D graphs or by 3D animation. The inflation and contraction of the entity nodes represents the accumulation and degradation of reactants in the pathway. In this way, complex biological processes with multiple components can be modelled.

### The interferon-β signalling network-an oscillating feedback control system

Biological systems display a variety of dynamic behaviours ranging from stable steady states to oscillations. Oscillations in protein concentrations or gene expression levels are commonly associated with the presence of negative feedback loop(s) in the regulatory network [7]. Based on the analyses performed using this system many factors can affect the amplitude, frequency and stability of oscillations. For instance, the time-delay (path length) between token input and inhibitor, the type of inhibition edge used (competitive or non-competitive), the number of inhibitors present and their halflife will allow affect how a system containing a negative feedback loop will operate in practice.

As an example a simple model of the interferon-β signalling pathway is presented. Interferon-β is a cytokine released by immune cells in response to pathogens. It acts as an autocrine and paracrine signalling system that triggers the activation of host defence systems. In the form provided the model operates as a damped oscillator [8], where low-dose IFN stimulation yields oscillations of lesser amplitude that are damped faster than those induced at a high-dose (Figure 5). To reproduce these observations users can modify the token input or topology of the example file provided (**Supplementary Data 3**).

## BOX 5: Glossary

- **Arc**: In Petri nets, arcs are directional edges that connect a place to a transition or vice versa, (but never between places or between transitions). Arcs are represented as **edges** in mEPN notation.
- **Edge**: In mEPN notation, edges are arrows used to connect entity and process nodes to indicate an interaction and also the interaction direction. The Petri net equivalent is an **arc**.
- **Entity node**: In mEPN notation, entity nodes are glyphs that represent model components such as molecules or genes. The Petri net equivalent is a **place**.
- **Glyph**: a visual representation of an entity, process or transition node.
- **GraphML**: a XML-based file format used to describe the structural properties of a graph.
- **Layout**: the way in which nodes and edges are set out in 2D or 3D space (physical topology).
- **Model**: the visual representation of a network.
- **mEPN**: modified Edinburgh Pathway Notation scheme is a graphical notation system based on the concepts of the process diagram.
- **Network**: a number of interconnected entities.
- **Notation scheme**: a system of symbols used to represent diverse biological entities and interactions in semantically and visually unambiguous manner.
- **Parameterization**: defining the initial state of the system by the placement of tokens.
- **Petri net**: is a directed bipartite graph that alternates places (entities) and transitions (events that occur).
- **Place**: In Petri nets, places represent possible states of the system. They are represented as entity nodes in mEPN notation.
- **Process diagram**: is a diagram used to give an abstract description of the behaviour of a system that represents changes of the state of each molecule using arrows that indicate **transition**.
- **Process node**: In mEPN notation, process nodes are glyphs that represent and define interactions between entities or the transition of an entity from one state to another. The Petri net equivalent is a **transition**.
- **SPN**: Signalling Petri net, a non-parametric model for characterizing the dynamics of signal flow through a signalling network using token distribution.
- **Tokens**: Tokens represent quantitative information that is introduced and distributed through a Petri net
- **Topology**: arrangement of various elements of the pathway (nodes, edges, etc) that illustrates how information flows within the network.
- **Transition**: In Petri nets, transitions are events or actions. They are represented as process nodes in mEPN notation.

## Supplementary Material

**Supplementary Data 1**: mEPN palette GraphML file

**Supplementary Data 2**: Interferon-β components GraphML file

**Supplementary Data 3**: Interferon-β pathway SPN ready model GraphML file

**Supplementary Data 4**: Video of Interferon-β pathway simulation

**Supplementary Data 5**: Example 1 model

All supplementary files can also be downloaded from:www.virtuallyimmune.com.

## Acknowledgements

We are grateful to Paul Digard and David Hume for helpful discussions and advice. We thank Athanasios Theocharidis and Derek Wright for BioLayout *Express*^3D^ development. We also thank the BBSRC who are currently funding the development of the program (BB / F003722/1) together with the Wellcome Trust (GR077040RP) who previously provided support.

## Author contributions

A.L., L.O., M.E.P., L.B.S. and T.C.F. wrote and edited the manuscript. T.A. and D.W. assisted with development of BioLayout *Express*^3D^ and the VirtuallyImmune website. T.C.F. supervised the project.

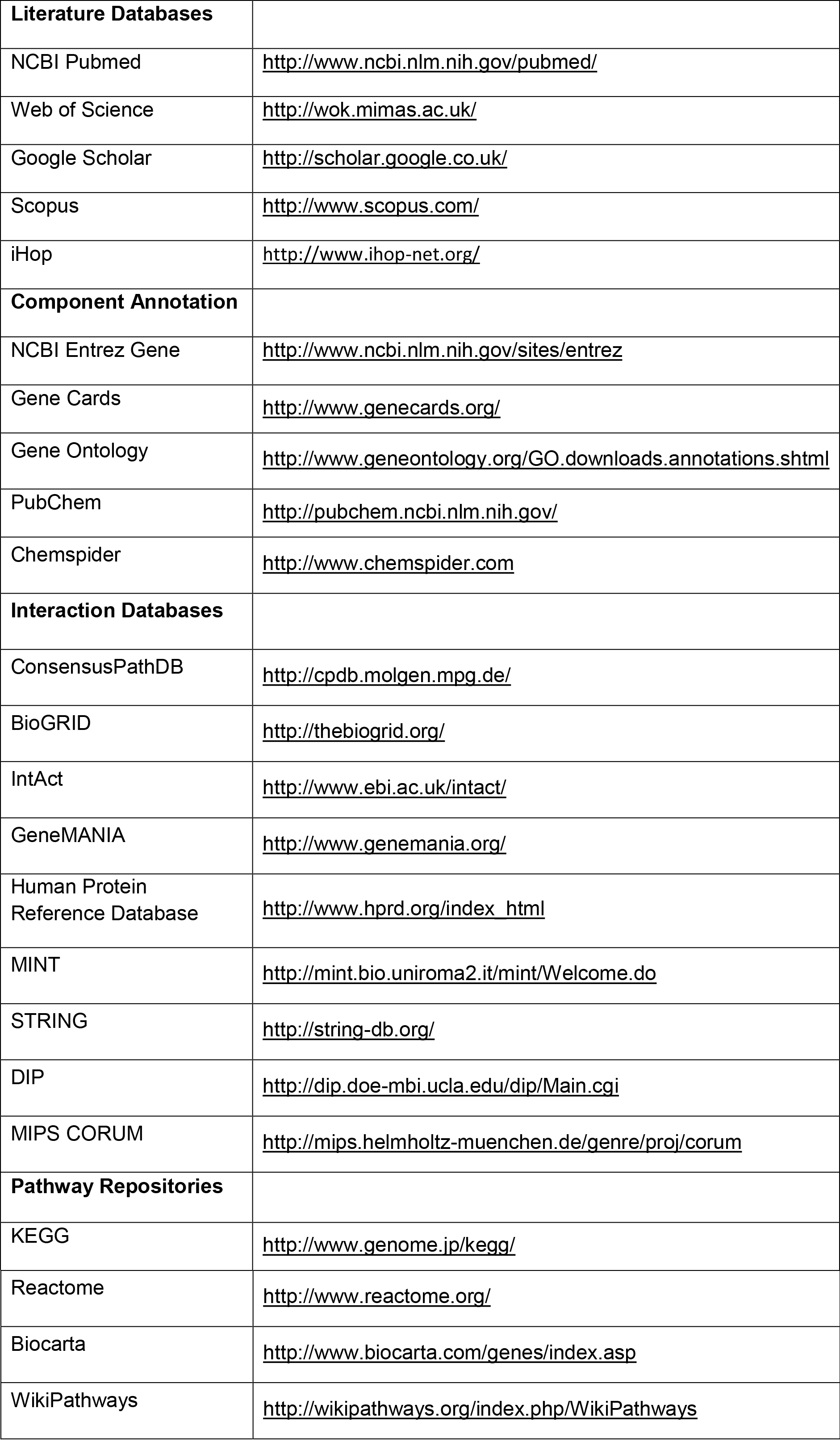
TROUBLESHOOTING

